# Spatially constrained tumour growth affects the patterns of clonal selection and neutral drift in cancer genomic data

**DOI:** 10.1101/544536

**Authors:** Kate Chkhaidze, Timon Heide, Benjamin Werner, Marc J. Williams, Weini Huang, Giulio Caravagna, Trevor A. Graham, Andrea Sottoriva

## Abstract

Quantification of the effect of spatial tumour sampling on the patterns of mutations detected in next-generation sequencing data is largely lacking. Here we use a spatial stochastic cellular automaton model of tumour growth that accounts for somatic mutations, selection, drift and spatial constrains, to simulate multi-region sequencing data derived from spatial sampling of a neoplasm. We show that the spatial structure of a solid cancer has a major impact on the detection of clonal selection and genetic drift from bulk sequencing data and single-cell sequencing data. Our results indicate that spatial constrains can introduce significant sampling biases when performing multi-region bulk sampling and that such bias becomes a major confounding factor for the measurement of the evolutionary dynamics of human tumours. We present a statistical inference framework that takes into account the spatial effects of a growing tumour and allows inferring the evolutionary dynamics from patient genomic data. Our analysis shows that measuring cancer evolution using next-generation sequencing while accounting for the numerous confounding factors requires a mechanistic model-based approach that captures the sources of noise in the data.

**Summary:** Sequencing the DNA of cancer cells from human tumours has become one of the main tools to study cancer biology. However, sequencing data are complex and often difficult to interpret. In particular, the way in which the tissue is sampled and the data are collected, impact the interpretation of the results significantly. We argue that understanding cancer genomic data requires mathematical models and computer simulations that tell us what we expect the data to look like, with the aim of understanding the impact of confounding factors and biases in the data generation step. In this study, we develop a spatial simulation of tumour growth that also simulates the data generation process, and demonstrate that biases in the sampling step and current technological limitations severely impact the interpretation of the results. We then provide a statistical framework that can be used to overcome these biases and more robustly measure aspects of the biology of tumours from the data.

## Introduction

Cancer is an evolutionary process [1] fuelled by genomic instability and intra-tumour heterogeneity (ITH) [2]. ITH leads to therapy resistance, arguably the biggest problem in cancer treatment today [3]. Recently, seminal studies have attempted to quantify ITH by either looking at subclonal mutations in deep sequencing data from single bulk samples [4], or by taking multiple samples of the same tumour, the so-called multi-region sequencing approach [5,6]. Phylogenetic approaches are then used to reconstruct the ancestral history of cancer cell lineages [7]. However, one important difference between phylogenetic analyses in cancer and classical phylogenetic analyses of species is that each cancer sample is not a single individual, but a mixture of different cancer cell subpopulations and non-cancer cells [8]. Even when multiple samples are collected, each sample is a bulk of (many) different cells.

The problem is usually tackled by performing a subclonal deconvolution of the samples to separate the different subpopulations [4,9]. However, these approaches do not account for the spatio-temporal dynamics that generated the data. To study the spatio-temporal evolutionary dynamics of individual tumours, mathematical and computational models of evolutionary processes are widely employed [10-12]. Many of these models are rooted in theoretical populating genetics, a field that quantifies the evolution of alleles in populations and that is central to the modern evolutionary synthesis [13]. More recently, complex spatial models have also been used [14-19]. However, seldom have mathematical and computational models of cancer evolution been directly connected to next-generation sequencing data from human tumours. Recent work from us and others has shown that combining theoretical modelling and cancer genomic data allows for measurement of fundamental properties of the tumour evolutionary process *in vivo*, effective mutation rates in tumours as well as the strength and onset of sublconal selection events [19-21].

Here, we study how spatial considerations impact our ability to infer cancer evolutionary dynamics. We combine explicit spatial evolutionary modelling with synthetic generation of multi-region bulk and single-cell data, thus providing a generative framework in which we know the evolutionary trajectories of all cells in a tumour and can examine the genomic patterns that emerge from the sampling experiment. We show that spatial constrains, stochastic spatial growth and sampling biases can have unexpected effects that confound both the interpretation and inference of the perceived evolutionary dynamics from cancer sequencing data. We also present a statistical inference framework that is able to account for some of these confounding factors and recover aspects of the cancer evolutionary dynamics from various types of multi-region sequencing data as well as single-cell data.

## Results

### Simulating spatial tumour growth, tumour sampling and sequencing data generation

Here we develop and analyse a stochastic cellular automaton modelling spatial tumour growth that incorporates cell division, cell death, random mutations and clonal selection (Material and Methods). Each tumour simulation starts with a single ‘transformed’ cell in the centre of either a 2D or 3D lattice, and we model the resulting expansion of this first cancer cell. All events, such as cell proliferation, death, mutation and selection are modelled according to a Gillespie algorithm [22].

In our model we account for different spatial constraints that are parameterised within our simulation. In order for a cell to divide, a new empty space for its progeny is required within the e.g. 8 neighbouring cells if we consider a 2D grid with Moore neighbourhood. If no empty space is present a cell can generate a new space by pushing neighbouring cells outwards (choosing a random direction of the push). In this scenario, the growth is *homogeneous*, where all cells in the neoplasm can divide (Figure 1A). Because all cells in the tumour can divide, this scenario leads to an overall exponential expansion (Figure S1A). At some point during the simulation (Figure 1A-D), within the original tumour population (blue cells), we introduce a new mutant (a new subclone – red cells) which may or may not have a selective advantage. In the case of a neutral subclone (no selective advantage), the mutant cells divide exactly as all the other cells (Figure 1A). We note that in this case, colouring a new subclone in red at a certain point during neutral growth is arbitrary, and equivalent to the marking of a lineage by a random neutral (passenger) mutations. In the case where the subclone has a fitness advantage, the mutant will, on average, grow more rapidly compared to the parental background clone, thus increasing in relative proportion in space over time (Figure 1B and S1B).

**Figure 1.**
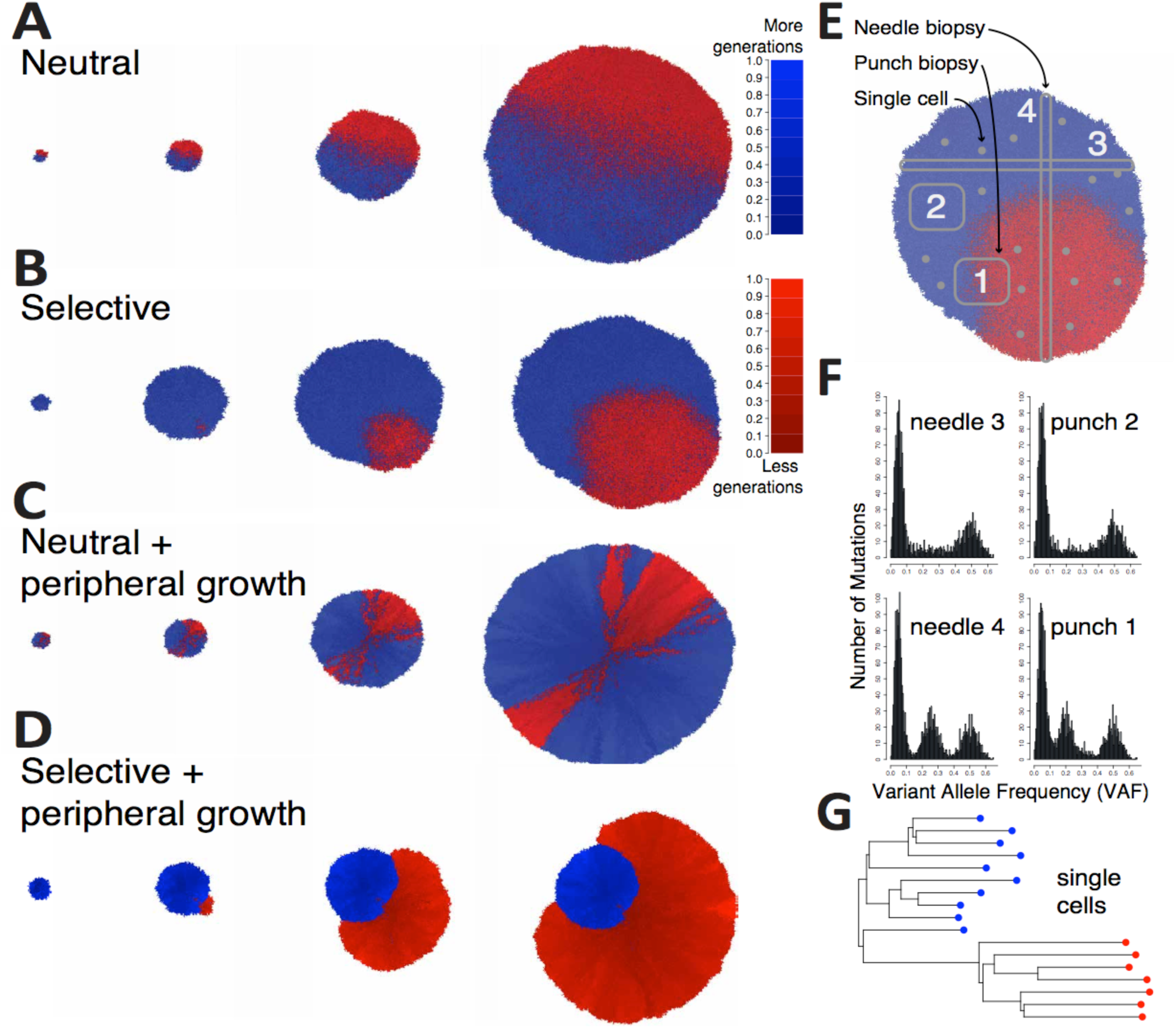
A spatial tumour growth model that simulates sequencing data. In our model we introduce a mutant at a given time t (blue = background clone; red = mutant subclone; shade is proportional to the number of generations the cell has gone through). **(A)** The new mutant subclone can have no fitness advantage (mutation is a passenger), giving rise to a neutrally growing neoplasm, or **(B)** have a fitness advantage s>0 with respect to the background population (mutation is a driver), giving rise to differential selection in the tumour population. In addition, cells accumulate unique passenger mutations during each cell division. **(C)** In many tumours, it is likely that only cells close to the tumour border are able to proliferate due to the abundance of resources and space. We simulate this in our model as boundary driven growth, which gives rise to complex radial patterns. **(D)** When boundary driven growth is combined with selection, spatial effect can either amplify the growth of the new subclone, as in this exemplary case, or even decrease the effects of selection if the subclone, by chance, gets imprisoned behind the growing front. **(E)** In our simulation we also model the raising and spread of point mutations in the genome of cancer cells (all passengers and, when subclone is selective, one additional driver). We can simulate the sampling of punch biopsies (squares), needle biopsies (thin stripes) and single cells. **(F)** By adding to the simulated mutations, the noise and measurement errors of next-generation sequencing, we can generate synthetic realistic variant allele frequency distributions. **(G)** Single-cell data can also be simulated, in this case clearly showing the presence of a selected subclone demonstrated by the clade of “red” cells with a recent common ancestor.

We also model ‘boundary driven’ growth, where only cells that are sufficiently close to the border of the tumour can proliferate. Other cells may remain ‘imprisoned’ in the centre of the tumour unable to proliferate because of the lack of empty space around them. Boundary-driven growth is a phenomenon routinely observed in real tumours when Ki67 proliferation staining are applied [23]. The magnitude of this effect is controlled in our simulation with the parameter *a*, which takes into account cell location and defines the probability that a cell will push neighbouring cells to create empty spots (see Materials and Methods). Boundary driven growth leads to a polynomial expansion (Figure S1C) of tumour cells over time. Importantly, in both the case of neutral mutants (Figure 1C) and selected mutants (Figure 1D), the spatial distribution of mutant cells is strongly affected by the spatial constraints.

At each division, a cell has a certain probability to acquire additional somatic mutations, modelled with a Poisson distribution, with mean *u*, in line with many other previous models [12,20,21,24,25]. Notably, u is the average number of new somatic mutations per division for the whole genome of a single cell. We assume that both daughter cells can acquire mutations, mutations are unique (infinite site model) and neglect back mutations (infinite allele model). Finally, in our model the large majority of mutations are assumed to be passengers (neutral), with a few driver alterations allowing for subclonal fitness advantages (e.g. subclonal populations in Figure 1B and D). This is plausible, as large-scale genomic sequencing studies indicate that in any given tumour the number of driver events is small, and the number of passengers many orders of magnitude larger [24,26].

Importantly, our spatial model of tumour growth allows for the simulation of tissue sampling and genomic data generation. For instance, we can simulate the collection of punch biopsies where spatially localised chunks of tumour are collected (Figure 1E). We can also simulate needle biopsies, where a long and thin piece of tissue is sampled (Figure 1E). We can then simulate the genomic data generation process starting from the cells in the sample and the identification of somatic mutations. For example, we can simulate the sequencing at a given coverage using Binomial sampling of the alleles, the limits of low frequency mutation detection (e.g. minimum number of reads with a variant, minimum coverage), as well as non-uniformity of coverage leading to over-dispersion of the variant allele frequency (VAF) of detected mutations. This allows generating realistic data from simulated tumours, e.g. in the case of the simulation of a diploid tumour with one selected subclone in Figure 1E, all needles and punch biopsies contained clonal mutations, shown as a cluster of variants around VAF=0.5, and in the case of punch biopsy 1 and needle biopsy 4, also a subclonal cluster representing the growing subclone (Figure 1F).

We previously showed, using a non-spatial stochastic branching process model of tumour growth, that assuming a well-mixed population and exponential growth, the expected VAF distribution of subclonal mutations in cancer under neutral growth follows a power-law with a 1/f^2^ scaling behaviour, where f is the variant allele frequency of subclonal mutations [20]. This has been previously demonstrated to be the scaling solution of the fully stochastic Luria-Delbruck model [27-29]. The 1/f-like neutral subclonal tail can be observed in needle 3 and punch 2 in Figure 1F. In the presence of subclonal selection we expect to observe an additional subclonal ‘cluster’ of mutations all at the same frequency [4], that are the passenger mutations hitchhiking in the expanding clone (as we previously described [30]). This is exemplified in needle 4 and punch 1 in Figure 1F. We note that a 1/f-like tail remains in the VAF frequency spectrum of all samples, as a consequence of within-clone neutral dynamics that remain on-going throughout the tumour’s growth [30]. Furthermore, our framework allows simulating single-cell data. For example, in Figure 1B we sample individual cells at random and simulate single-cell whole-genome sequencing (Figure 1G).

### Spatial effects on bulk sequencing data: drift, draft and sampling bias

For each representative simulation of spatial constrains in Figure 1, we modelled the sampling of 6 punch biopsies (small square regions), 2 needle biopsies (long and thin regions), as well as hypothetically sampling the whole tumour. From each sample, we simulated the generation of 100x depth whole-genome data (see Material and Methods for details about the noise model). Figure 2A shows the variant allele frequency (VAF) distributions of samples from the neutral homogeneous growth case in Figure 1A, with clonal mutations (truncal) in grey, subclonal mutations exclusive to the parental background clone in light blue and subclonal mutations within the mutant in pink. As discussed before, all samples show the characteristic 1/f^2^distribution corresponding to neutral evolutionary dynamics [20], as one would expect theoretically [27]. The proposed R^2^-based frequentist test for neutrality we previously proposed [20] is reported on top of each VAF plot and shows that even in the presence of a spatial structure, homogeneous (exponential) neutral growth follows a 1/f^2^ distribution (Figure 2A-i to 2A-iv). As we have shown previously, it is possible to recover the effective mutation rate (mutation rate per cell doubling) from the ∼1/f^2^ neutral tail, which in this case without cell death was 10 mutations per division (∼10^−9^ mutations/bp/division). This was correctly recovered in all samples from Figure 2A (recovered mutation rate reported in each plot as *u*).

**Figure 2.**
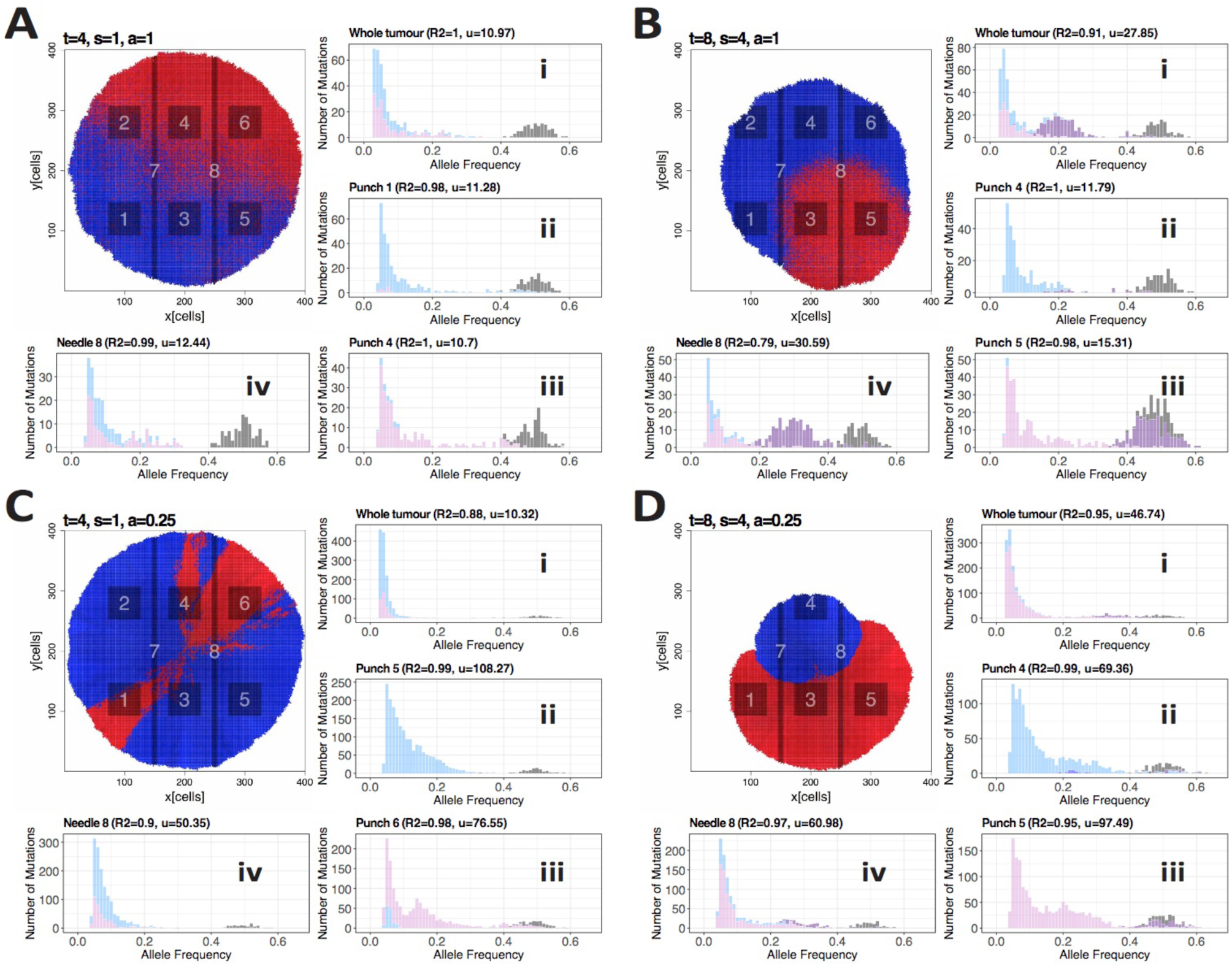
Variant allele frequency distributions of punch and needle biopsies from representative scenarios. **(A)** In the illustrative example of neutral homogeneous growth, a neutral mutant was introduced at generation time t=4 with a selection coefficient of s=0 (neutral) and homogeneous growth (a=1). The mutation rate was u=10. Tumour was simulated until ∼100,000 cells. From the final tumour, we sampled 6 punch biopsies (1-6), 2 needle biopsies (7-8) and a “whole-tumour” sample, and simulated 100× whole-genome sequencing data. VAF distributions of each sample are shown (i-iv). Colours: grey = clonal mutations; light blue = mutations subclonal in the background clone; light red = mutations subclonal in the mutant clone). **(B)** In this case, a differentially selected subclone with s=3 was introduced at time t=8 in a homogeneous growth scenario (a=1) and u=10. Final population size was ∼80,000 cells. In those samples where both the background and the mutant subclone were present (B-i and B-iv), the VAF distribution showed evidence of subclonal selection, with a subclonal cluster (purple) generated by mutations in the selected subclone that hitchhiked to high frequency due to selection. **(C)** In the case of neutral boundary driven growth, a new (neutral) mutant was introduced at t=4 with s=1 and boundary driven growth parameter a=0.025. Even though the tumour grew neutrally, the spatial effects of boundary driven growth led to deviations from the neutral expected null (which was derived based on homogeneous growth). Moreover, false “clusters” in the VAF spectrum are evident in C-3, where sampling bias produced an over-representation of a lineage that was not due to selection. **(D)** Boundary driven growth with selection (mutant introduced at t=8 with s=2 and a=0.025) produced even more complex patterns of drift and sampling bias.

In the case of homogeneous growth with subclonal selection (Figure 2B), neutrality could be rejected based on the subclonal tail in those samples containing a mix of the background clone and the new subclone (Figure 2B-i and 2B-iv, see subclonal cluster in purple). Specifically, needle 4 and punch 1 showed the expected signature of selection, with a subclonal cluster a consequence of the over-representation of passenger mutations in the expanded clone [4,21]. The 1/f^2^-like tail resulting from the within-clone accumulation of passenger mutations remains in the frequency spectrum [21]. Specifically, in the plots in Figure 2B we report the mutations that were present in the first subclone cell in purple. Those are mutations that increased in frequency by hitchhiking on the selected mutant. Importantly, we note that these mutations are not exclusive to the subclone but are also found in other lineages (e.g. in the ‘cousins’ of the selected subclone). Selection could not be detected in other spatially-distinct samples from the same tumour, as these did not contain differentially selected populations, and either captured only the background clone (blue) or only the selected mutant (red) (Figure 2B-2 and 2B-3).

This initial spatial analysis produced similar results to our previous well-mixed non-spatial models [20,21]. We next investigated the effect of boundary driven growth. Here, only cells close to the borders grow, leaving other cells ‘imprisoned’ inside the tumour mass (see Material and Methods for details), a pattern of *gene surfing*, or sometimes called *genetic draft* emerges, causing radial patterns of cells growing only at the front of the growing wave (Figure 2C). This has been previously documented both theoretically and experimentally in bacteria [31], as well as cancer [15,16,32]. Because the population is no longer homogeneously distributed, this can lead to significant spatial sampling bias, causing over- or under-representation of mutations in the VAF distributions solely due to spatial effects and not because of selection. This causes deviations from the neutral expectation of the mutant allele distributions that risk being wrongly interpreted as the consequence of on-going subclonal selection, as in Figure 2C. In this scenario, we know that subclonal clusters are *not* genetically distinct differentially selected subclones, but the over-representation of alleles compared to the neutral expectation is solely induced by the spatial structure. Furthermore, even when we observe distributions that appear to follow the neutral expectation (high R^2^), boundary driven growth results in much higher mutational loads than would be expected in the well mixed case. Here our inferred mutation rates are up to 10 times higher than the ground truth. This can be observed from Figure S2, where we sample each representative tumour from centre towards periphery by taking samples along concentric circles of the tumours and compare the mutational loads of the samples. A similar phenomenon is also observed in species evolution [33].

If we combine boundary driven growth and subclonal selection the situation is further complicated: selective effects are now modulated by spatial constraints. In some cases, the selected mutant emerges and remains directly at the front of tumour growth. In this scenario the outgrowth caused by its selective advantage is amplified further just because it occurred at the growing front (Figure 2D). In other cases, the selected mutant may, by chance, remain ‘imprisoned’ within the tumour (assuming the mechanism of selective advantage is unable to overcome this spatial entrapment) and stops proliferating despite its selective advantage (Figure S3). In both these cases, further sampling biases occur. In the case of punch 5 for example (Figure 2D-3), where the new subclone is fixed (clone fraction=100%), there is an overrepresentation of a cluster of mutations that is only due to spatial drift and not selection. These dynamics are recapitulated in larger cohorts of simulated tumours with the same parameters (Figure S4).

We then looked at the pairwise VAF distributions between samples. The amount of subclonal mutations scattered through the frequency spectrum (Figure 3A-D) and the number of false subclonal clusters due to sampling bias and spatial drift was striking (e.g. Figure 3D). As per ground truth, only the purple mutations shoud show a subclonal clustering pattern (e.g. Figure 3B, punch 1). We found that scattered variants were mostly due to the effect of neutral lineages spreading in space, and then be subsampled in different ways in each tumour region. In the case of boundary driven growth, sampling bias produces evident clusters that do not correspond to differently selected subclones in the tumour. This makes the reconstruction of the true clonal phylogeny and its evolutionary interpretation very problematic.

**Figure 3.**
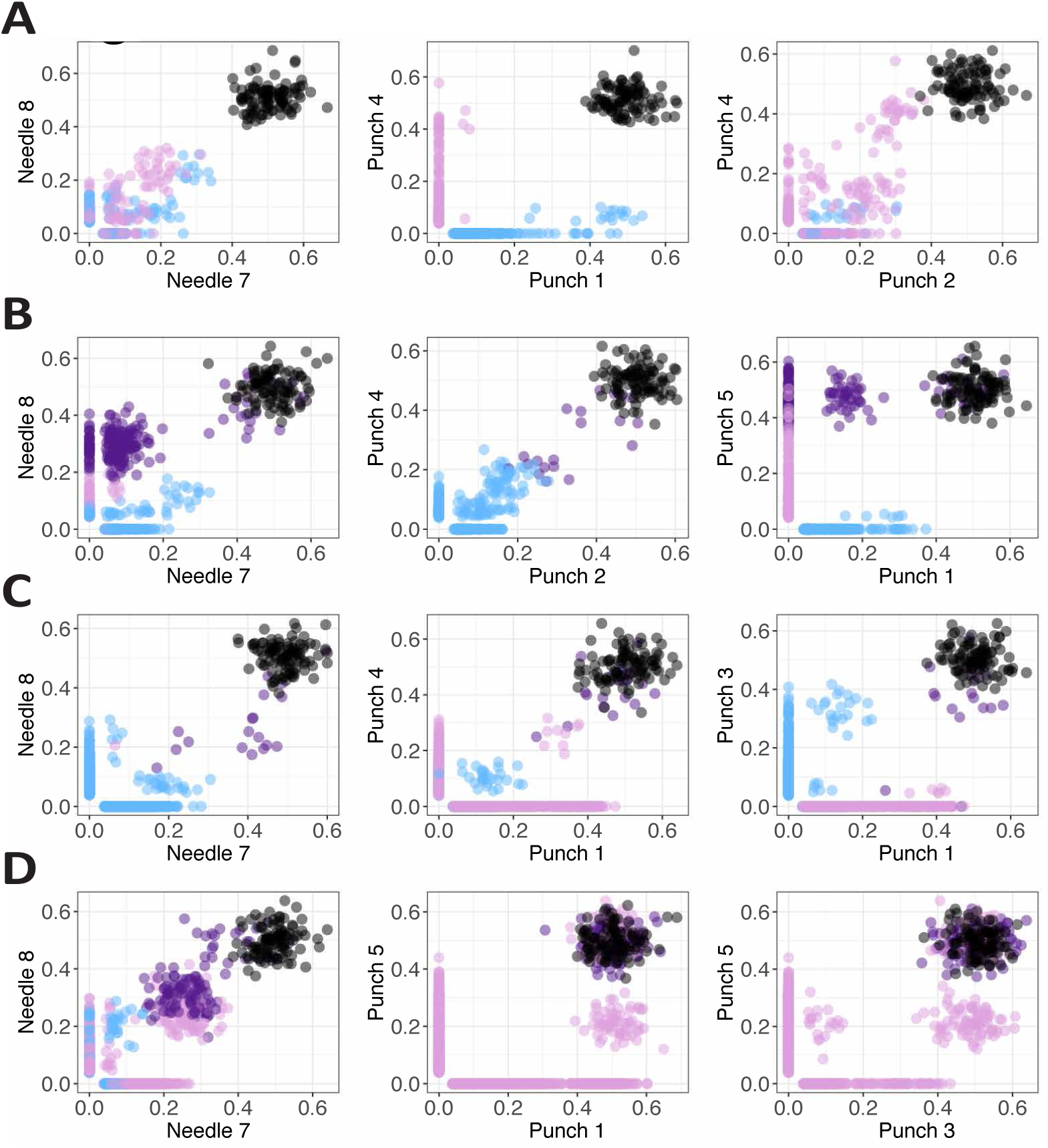
Sample vs sample scatterplots of mutations for each case. For each of the representative cases: **(A)** neutral homogeneous, **(B)** selective homogeneous, **(C)** neutral boundary driven, **(D)** selective boundary driven, we report the scatterplots of somatic mutations in selected samples. Clearly, the presence of passenger subclonal mutations in the neutral tail of growing clones that spread in space as the tumour grows produces scattered variants (e.g. A). Even more striking is the formation of false subclonal clusters of mutations particularly in the presence of boundary driven growth (e.g. C, D) where some lineages are over-represented not because of differential selection, but because of sampling bias and spatial drift.

### Spatial effects on single-cell sequencing

Most of the confounding factors we have described so far result from the limitations of bulk sequencing, where the genomes of many cells are convolved within samples. Single-cell sequencing does not suffer from this particular limitation and promises a big leap for cancer evolutionary analysis [34].

To examine the effect of single cell sequencing, we simulated the whole-genome sequencing of 10 single cells taken at random from the tumour and reconstructed the phylogenetic relationship between these cells (Figure 4). For the neutral cases (Figure 4A and C), the patterns are consistent with a typical ‘balanced’ neutral tree, wherein all lineages contribute roughly equally to the final cell populations. In cases with subclone selection (Figure 4B-i and D-i), the selected subclones are over-represented on the tree (analogous to the behaviour seen in VAF distributions). The pattern is even clearer if we sample 400 single cells and performed WGS (Figure 4B-ii and D-ii). We note that if we use randomly sampled single cell sequencing and plot the site frequency spectrum (frequency distribution of mutations within the population of sampled cells) we recapitulate the VAF distribution (Figure S5). This is because the site-frequency spectra derived from single-cell sequencing data corresponds to a VAF distribution.

**Figure 4.**
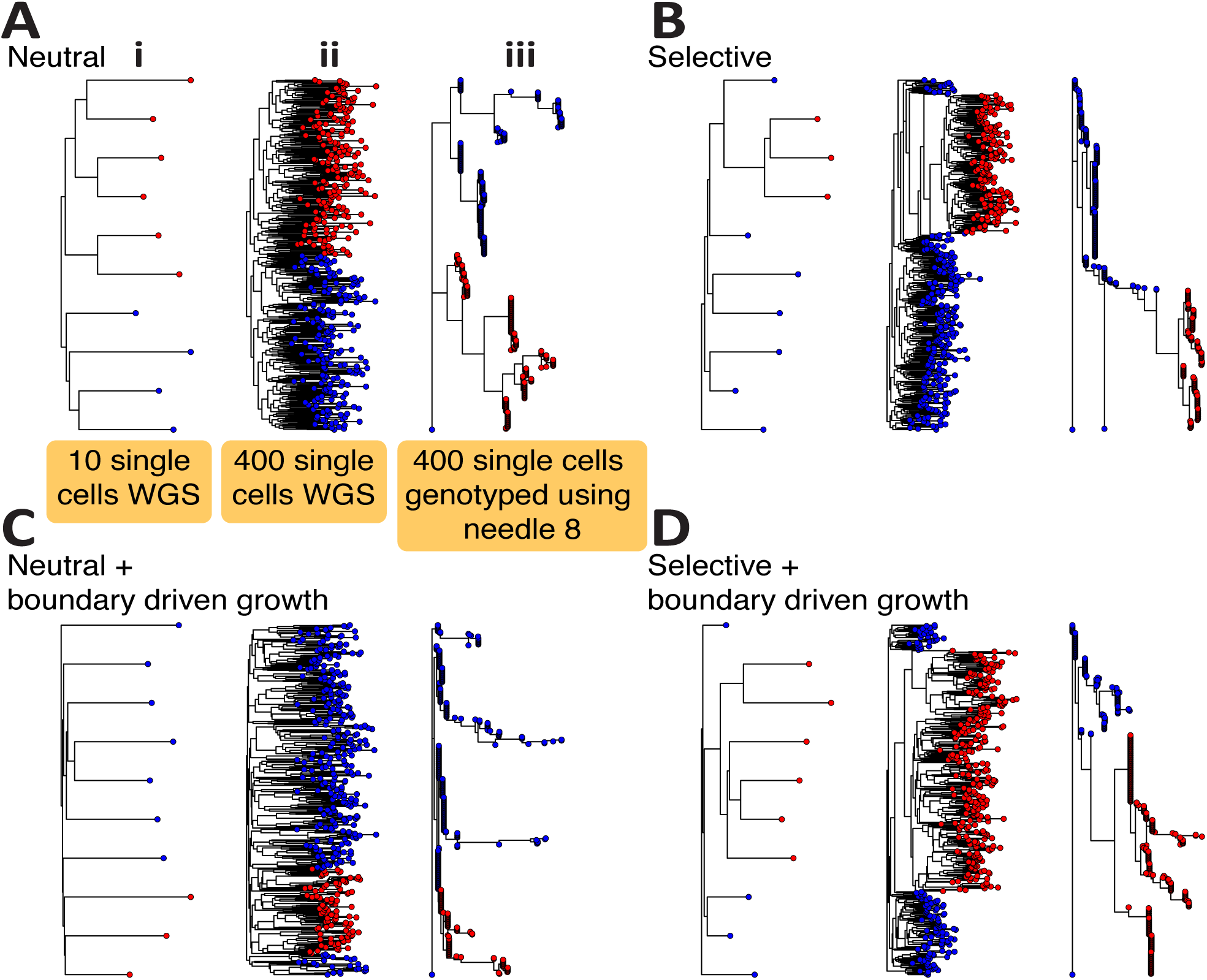
Single-cell sequencing data from spatial tumour simulations. **(A)** From each representative scenario we sampled 10 single-cells at random (i) as well as 400 single-cells at random (ii) and performed synthetic whole-genome sequencing. In all cases, single-cell sequencing significantly reduces the sampling bias that we found in bulk samples and the only overrepresented lineages were due to selection (B, D). However, due to the currently high error rate of single-cell sequencing, several studies rely on single-cell genotyping using mutations found in bulks. We simulated this by genotyping on 400 single-cells the mutations found at VAF>5% in needle biopsy 8 of each tumour (iii). The resulting trees are very hard to interpret in terms of the clonal phylogeny due to the bias in the selection of variants to genotype.

However, as whole-genome mutational profiling of single cells is still difficult due to allele dropout [35], often single-cell genotyping has to be performed instead [36]. For instance, a bulk sample is sequenced and all mutation in that bulk sample are then tested in single cells for presence/absence of the mutation. Integrating bulk sequencing with single-cell information is extremely powerful [37], but requires careful interpretation of the resulting data. In Figure 4A-3 we show that this approach, although informative, can lead to very distorted phylogenetic trees where branch lengths are heavily biased by the initial choice of mutations to be assayed, and consequently the signature of selection vs neutrality is not readily identifiable from these data alone.

Moreover, significant sampling bias is still apparent for single-cell sequencing when individual cells are not sampled uniformly at random from the whole tumour, but instead isolated in ‘clumps’ from different bulk samples. In Figure 5 we have simulated the collection of 4 single cells from each of the 6 punch biopsies in Figure 2. The trees are quite different from Figure 4 and moreover, it is interesting to see how the underlying patterns of growth are reflected in the mixing of cells from different bulks. For instance, homogeneous growth leads to very high intermixing of cells in different bulks, whereas boundary driven growth tends to spatially segregate in bulks. We have seen these patterns in carcinomas vs adenomas, where carcinomas were characterised by clonal intermixing, but adenomas were not [17]. Similar patterns of intermixing have been demonstrated recently using single-cell seeded organoid sequencing [38].

**Figure 5.**
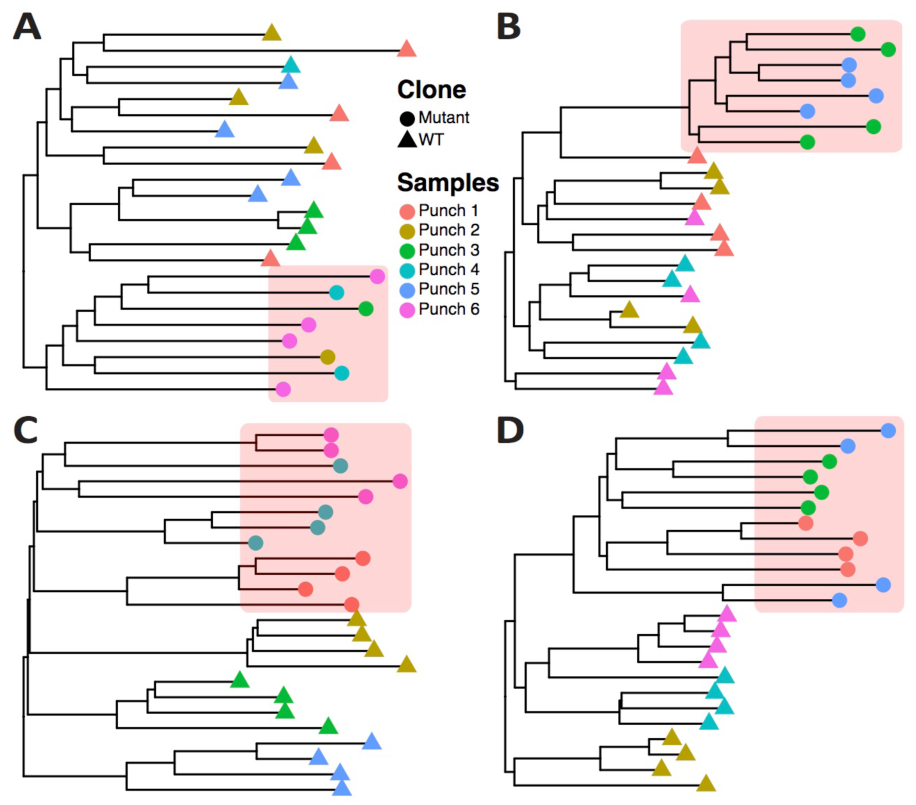
Biases of single-cell sequencing when cells are taken from spatially separated bulk samples. Whereas taking N random cells from a tumour highly reduces sampling bias, this is often not how single-cell from neoplasms are sampled. Often first small chunks of the tumour are dissected and then single-cells are isolated from those. **(A)** neutral homogeneous, **(B)** selective homogeneous, **(C)** neutral boundary driven, **(D)** selective boundary driven. For each of our representative examples, we simulated this type of sampling and show how this impacts severely on the phylogenetic tree and patterns of clonal intermixing. In particular, it alters the detected phylogenetic relationship of the cells because, since groups of cells come from spatially segregated regions, those appear more closely related than expected by chance. This is an important source of sampling bias that needs to be considered when analysing single-cell phylogenies. Cells coming from the ‘red’ mutant subclone are highlighted in the red shaded box.

### Resolving spatial effects with inference

The spatial effects of drift and sampling bias one can observe are remarkable and represent a major challenge for the correct subclonal reconstruction of tumours growing in three-dimensional space. Due to the inherent complexity, analytical solutions to this problem that take space into the account remain challenging, although some attempts to tackle this difficult question are being undertaken [39]. However, understanding the complex impact of spatially growing cell populations on the actual genomic data requires an approach based on computational simulations.

Here we devise a statistical inference framework, similar in spirit to what we previously proposed for well mixed populations [21], that aims at recovering the evolutionary parameters of each individual tumour from the type of data we have discussed so far (see Material and Methods for details). We constructed a test-set of 30 synthetic tumours simulated with different parameters (see Table S1) and assessed the error in recovering the parameters used to generate these tumours after statistical inference with an Approxiate Bayesian Computation – Sampling Monte Carlo (ABC-SMC) approach [21,40-42]. We were particularly interested in comparing the accuracy provided by the different spatial sampling methods in recovery evolutionary dynamics. We studied three different sets of tumours. In the first set, we investigated parameter recovery in tumours with homogeneous (exponential) growth, with and without selection but with no cell death. In the second set, we added stochastic cell death as an additional factor. In the third set, we studied cases of boundary driven growth where we also examined our ability to recover the extent of the boundary driven growth factor (i.e. how close a cell has to be to the border to proliferate – modelled by the parameter *a*). In all three sets, we studied the differences in the ability to recover parameter if we used a single bulk sample of the whole tumour multi-region punch biopsies, multi-region needle biopsies or single-cell sequencing.

Not surprisingly, the scenario with exponential homogeneous growth without cell death was the one where the evolutionary parameters were the easiest to recover because spatial constrains were limited and the number of unknown parameters lowest (Figure 6A-C). In particular, the percentage-error in recovering the mutation rate *u* was particularly low, especially using single-cell sequencing (Figure 6A). The mean percent error of the parameters *t* (Gillespie time when a new mutant is introduced) and *s* (selective coefficient of the new mutant), in the case of homogeneous growth were also within 20% and overall agrees with our previous observations in well-mixed populations [21]. The presence of stochastic cell death, even within a homogeneously growing tumour, introduced significant spatial and sampling biases (spatial drift) that led to a higher error in the recovery of the parameters (Figure 6A-C). Furthermore, some of the evolutionary parameters become not separately identifiable (mutation and death rate). In this scenario, the best sampling strategies to recovery the death parameter *d* were single-cell sequencing or whole-tumour sequencing, reflecting the need to collect large population of cells for the correct estimation of this parameter (Figure 6D). Boundary driven growth also introduced significant biases that led to higher percent-error values in the recovered parameters (Figure 6A-C). Here, single-cell sequencing was best in recovering the boundary driven growth parameter *a* (Figure 6E). See Figure S4 for VAF distributions and Figure S6 for summary statistics from the simulations in Figure 6. The full posterior distributions of each parameter in each context is reported in Figure S7. Parameter dependency in the inference of *t* and *s* combinations is reported in Figure S8.

**Figure 6.**
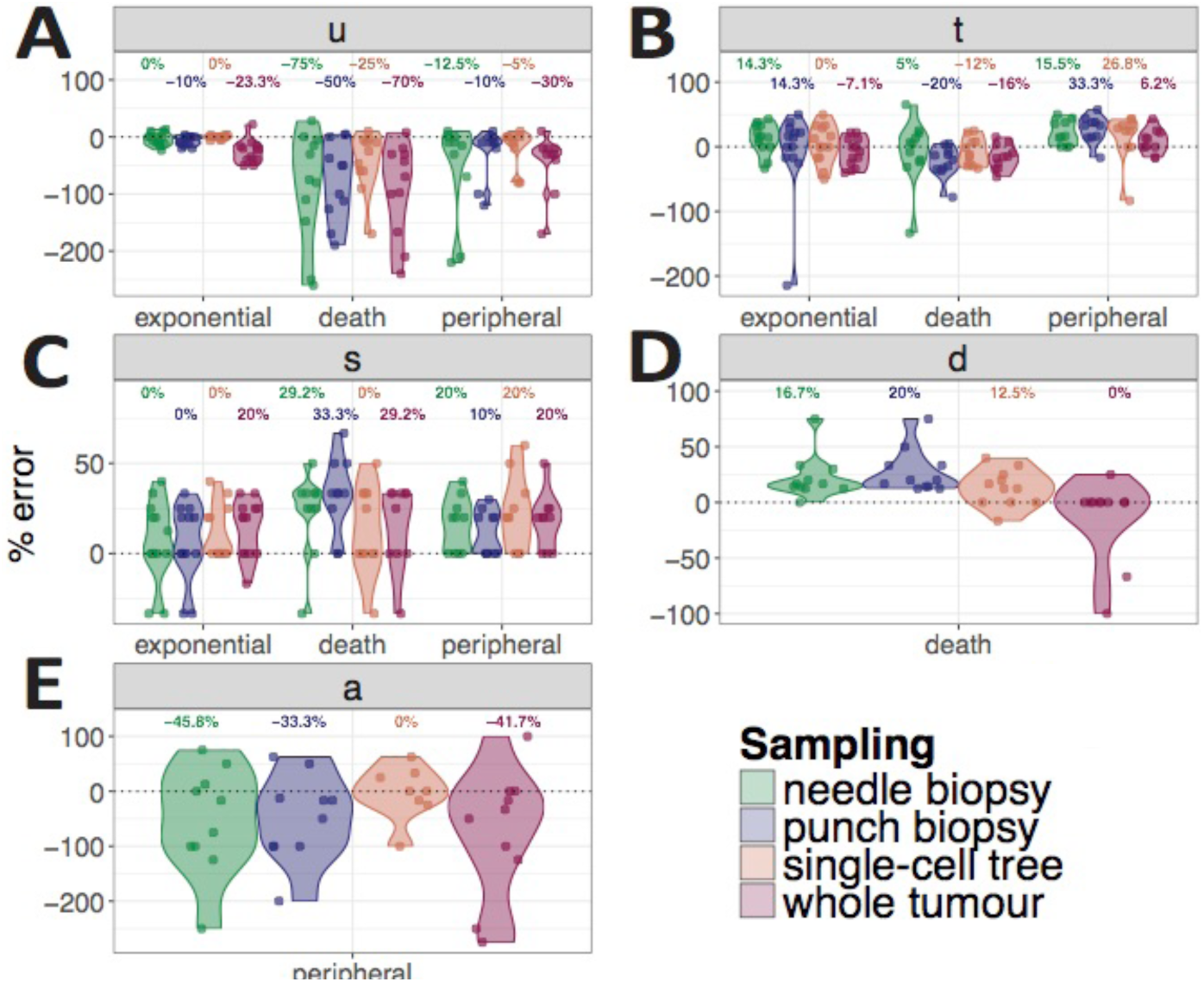
Statistical inference framework to recover evolutionary parameters. We combined our model with a statistical inference framework (Approximate Bayesian Computation – Sequential Monte Carlo) in order to infer the evolutionary parameters of selection and growth from the data. We tested this framework on 30 synthetic tumours for which we generated genomic data. We tested the ability to recovery parameters of 4 different sampling schemes: punch samples, needle biopsies, single cell phylogenetic trees and whole-tumour sampling (see Materials and Methods for details). We report the percentage error of the inference (true parameter value – inferred value based on the mode of the posterior probability) for each parameter and scenario. We investigated three different scenarios: **(A)** homogeneous growth without cell death, **(B)** homogeneous growth with stochastic cell death, and **(C)** boundary driven growth with different parameters values of a. Percent-error of the recovered parameters is reported for different sampling strategies. **(D)** For the homogeneous stochastic cell death scenario, we also report the error in recovering the death rate parameter d. **(E)** For the boundary driven growth scenario we report the error in recovering boundary driven growth parameter a.

## Discussion

It is now widely accepted that tumour growth is governed by evolutionary principles. Thus, recovering the evolutionary histories of tumours is essential to the understanding patient-specific tumour growth and treatment response. However, these inferences are necessarily based on limited information due to sampling biases, noise of known and unknown nature, lack of time resolved data amongst many others. Despite these limitations many approaches based on single or multi-region bulk or single cell sampling have been developed. Information from such studies is however often extracted based on clustering based analyses, without consideration of the confounding underlying influence of the cellular mechanics of tumour growth. Here we explicitly investigated spatial effects on the evolutionary interpretation of typical multi-region sequencing data of tumours. We found the effects more profound than we had anticipated. The effects of sampling biases and spatial distributions of spatially intermixed cell populations critically depend on the mode of tumour growth as well as the details of the underlying sampling and data creation procedure. Most surprisingly we could observe artefacts in the VAF distribution of some tumour samples that exactly appear as would positively selected subclonal populations but are solely due to the spatial distribution of cells. In our current understanding such artefacts are virtually indistinguishable from truly positively selected clonal subpopulations and cause a major challenge for the evolutionary interpretation of cancer genomic data.

We furthermore presented a Bayesian inference framework to recover evolutionary parameters from our spatial distributions. Again, we observe that our ability to precisely recover certain evolutionary parameters depend on the scenarios of tumour growth and spatial sampling strategies. However, we do believe that although complex, the situation is far from hopeless. More involved statistical frameworks based on first principals of tumour growth can help resolving some of the evolutionary parameters on an individualised patient basis. Importantly, careful spatial sampling and single-cell sequencing can mitigate some of the confounding issues. We do acknowledge that our model has some important limitations, such as the infinite allele assumption (which could be violated by copy number loss [35]). Also, for computational feasibility we only present 2D analyses and of a relatively limited number of cells with respect to the billions of cells present in a human tumour. However, our approach highlights the importance of spatial modelling of real data and the impact of confounding factor in our estimate and understanding of tumour evolution.

## Materials and Methods

### Details of the model

We developed a computational stochastic model of spatial tumour growth which incorporates different schemes of multi-region tissue sampling followed by imitating high-throughput sequencing procedures of data generation.

We consider tumour cells as asexually reproducing individuals that die and reproduce with certain pre-defined probabilities. If b is a birth rate for each cell and d a death rate, then exponentially expanding population can be modeled by the equation:

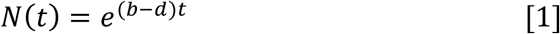

where N(t) is a population size at time t, and b-d is the net growth rate of the population. We assume that birth and death rates are constant over time, but growth rate can be stochastic throughout tumour development due to the stochasticity of event (death or birth) order and some spatial constrains that can limit or promote cell growth over time. For instance, we model spatial constraints with the aggression parameter – a, which is an outer radius of a growing tumour that defines its periphery where the cells are allowed to proliferate. In the limit of tumour expansion, when a is close to 1 (which is equivalent to 100% of tumour radius i.e. no spatial constrain), the growth is equivalent to exponential expansion. While when a is close to 0, the growth is extremely constrained as only the cells on the tumour periphery proliferate and hence the growth curve significantly diverges from the one given by a realization of equation [1].

In addition to the stochasticity of cell division, we have two other random events – mutation and selection, where the latter can change birth and/or death rates. _ We model somatic mutations that are acquired by each progenitor cells after cell division as a Poisson random variable – Pois(*u*), where u is the mean mutation rate. Thus after each cell division, a random set of new unique mutations are passed onto the offspring of the dividing cell. The majority of these mutations are passenger mutations and hence do not affect a cell’s phenotype. However, they enable us to trace cell lineages uniquely in the final tumour. In addition, we also allow for driver mutation ‘events’ that can lead to positive selection of a subpopulation of cancer cells: a driver event conveys a fitness advantage to that particular cell and its offspring and potentially allows the lineage to increase in frequency. Driver mutation events are scheduled in time at desired time-points, according to a parameter of the simulation.

To simulate tumour growth in space with these four stochastic events – birth, death, mutation and selection – we have used a modification of the Gillespie algorithm [22]. The simulation framework is as follows:

- **Initialization:** define a 2D/3D square grid that defines and constrains the possible spatial directions a tumour cell can grow to. Place the first tumour cell in the centre of the grid. Set initial time to 0. Until the simulation clock has reached a desired time T, repeat the following steps

1. Compute the reaction propensities according to the Gillespie algorithm. Each reaction event of birth (resp. death) has a functional form *f*(*x*) = *kx*; here *x* is the number of cells of type “x” (wild-type or mutant), and *k* is either the birth or death rate. The time of each event is obtained by sampling an exponential random variable with mean given by its propensity. The next event chosen is the one completing first (i.e., with smallest clock value, as in the so-called next reaction method [22]). Given the event, we increment time by its clock. Note that these time steps do not correspond to population doubling times i.e. generations; doubling times can be retrieved scaling time by a factor log(2).
  a. If the next event is a cell division, we use a heuristic method to place the 2 daughter cells on the grid. We first replace the parent cell with the first daughter, and search for a suitable position to place the second daughter cell. We use a Von Neumann neighbourhood and check if any of the 8 (in 2D grid) neighbouring spots of the parent cell are empty; if they are, we locate the second cell in one of those spots. Otherwise, with a probability determined by a parameter *a*, we push all cells along a randomly chosen direction until they hit the grid boundary, and place the second daughter at the first emptied spot. With this input parameter *a* we can model boundary driven versus exponential growth, as it represents the fraction of the radius of the growing tumour where cells are allowed to proliferate; that is, *a* =.2 creates a tumour periphery of the width equal to the 20% of the growing whole tumour in a given generation (hence when *a* = 1, periphery width is 100%, every cell can always push and divide, and the tumour grows exponentially). When a cell divides, we generate passenger mutations by drawing a number from Pois(*u*). These mutations will be assigned to both daughter cells.
  b. If the next event is cell death, we simply free the position allocated to the cell.
2. At the end of this step, we check if the clock is greater than the time of the next scheduled driver event; if it is, we convert a random wild type (WT) cell into a new mutant and increase its birth rate, or decrease its death rate. These mutant cells will therefore have a proliferative advantage. To quantify the effect, we define the fitness s as: 1 + *s* = (*birth*_*mutant* – *death*_*mutant*)/(*birth*_*wt* – *death*_*wt*).

#### Details of the data generation and error modelling

We also implemented schemes for bulk and single cell sampling. Bulk Samples are spatially separated tumour chunks ‘cut out’ from the whole tumour tissue. We model two different shapes:

1. Squares, which are referred to in the paper as punch samples,
2. and long thin rectangles that resemble a needle biopsy sampling strategy.

A bulk sample is a set of adjacent cells from the final tumour population. Each cell has its unique ID, a position on a grid and its list of somatic mutations. The mutations list contains the ancestral (to the given) cells mutations and the unique set of mutations acquired during the birth of the given cell. From the sampled cells (in a bulk) joined list of mutations we can construct the Variant Allele Frequency (VAF) distribution as in a real sequencing experiment.

To constructing a VAF distribution from a simulated bulk tumour sample, we mimic realistic next generation sequencing procedures. The simulation of high-throughput sequencing data requires introducing noise in sequencing coverage, as well as limits to the detectability of low frequency mutations. We proceed as follows:

1. We generate (dispersed) coverage values for the input mutations. We do that by sampling a coverage from a Poisson distribution *D*∼ *Poisson*(*λ* = *Z*) with mean *λ* equal to a desired sequencing depth *Z*;
2. Once we have sampled a depth value *k* for a mutation, we sample its frequency (number of reads with the variant allele) with a Binomial trial. We use *f*∼Binomial(*n, k*) where *n* is the proportion of cells carrying this mutation, out of all cells sampled in the simulated biopsy.

This procedure guarantees that the generated read counts reflect the proportions of mutations in the simulated tumour. To model detection limits for a mutation, we discard all mutations with *f* below a minimum values (in our case 5 with the desired coverage 100, which accounts for a ∼0.05 minimum VAF).

We also performed single cell sequencing taking either random single cells across the whole tumour population, or spatially structured biopsies where we uniformly sample single cells from within bulk samples. We used the obtained single cells to construct maximum parsimony phylogenetic trees and tried to assess the tree shapes that mirror the different scenarios of tumour evolution. In addition to single cell sequencing, we also model genotyping cells with a given list of mutations (targeted sequencing). To implement this, we took one of the bulk samples as reference genotype and check for presence of each individual mutation in a random set of 200 cells. Similarly, we use the obtained genotyped single cells to infer phylogenetic trees and check how much the genotyped trees differ from the single cells trees.

### Details of the ABC framework

In our complicated spatial model we do not have explicit formulas for the stationary probabilities of the stochastic process, and hence cannot derive a likelihood function. Thus, we have to use likelihood-free methods to compute posterior distribution of the parameters *θ*.

Here we use Approximate Bayesian Computation (ABC) [41,43] to infer the underlying evolutionary parameters of our model. ABC is based on the idea of scanning a large grid of plausible values for *θ*, and simulating the model with such parameterization. Outputs of the model are stored and compared to a predefined set of summary statistics that are initially evaluated on real data. We can then rank sets of parameters that lead to observed data which resembles our input. We can estimate a posterior distribution *p*(*θ*|*D*) for the model parameters *θ*, using the available data *D* and the prior for *θ*. This method is computationally intensive, and requires running several hundreds (ideally thousands or millions) simulations.

There are different approaches to implement ABC, the simplest is rejection-sampling. More advanced implementations such as ABC with Markov Chain Monte Carlo (MCMC) can result in significant increases in efficiency. In our paper we implemented a simple rejection-sampling algorithm first, and then added Monte Carlo simulation techniques to speed up the algorithmic convergence. The simple ABC rejection-sampling algorithm consists of the following steps:

1. Sample parameter vector *θ* from a prior distribution *p*(*θ*).
2. Run the model with the given parameter set and generate the corresponding dataset
3. Evaluate the distance between the simulated dataset and the target model data
4. If the distance is less than a desired threshold, accept the parameters.
5. Return to step 1 and repeat until *N* parameter values are accepted.

One of the most important factors that affect the ABC outcome is the number of simulations that one can afford to run, and the summary statistics chosen to evaluate the distance between a target and a simulated dataset. Summary statistics can be any quantitative measurement that captures the defining data characters and does not sacrifice too much information. As our distance metric, we used Euclidean and Wasserstein distances between summary statistics for different parameters as discussed below.

Wasserstein metric estimates distance between probability distributions by treating each distribution as a unit amount of dirt piled up on a given metric space and calculates the minimum cost required to convert one pile into another. If x and y are two vectors we want to evaluate the distance between, the metric considers them as locations on the real line of m deposits of mass 1/*m* for the vector x and n deposits of mass 1/*n* for y. Then after calculating the empirical distribution functions for both vectors: 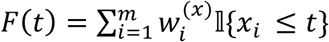 and 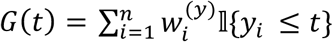 (for weights 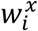 and 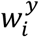 we took 1/*m* and 1/*n* respectively), the Wasserstein distance is defined by evaluating the following:

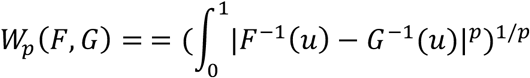

where we took p=1 for our analysis. We used the R package transport [ref (22)] to implement the distance calculation.

We used different summary statistics for each sampling scheme. For punch, needle biopsy or the whole tumour tissues – we used the VAF distribution to compute our summary statistics. For the whole tumour VAFs our ABC procedure was similar to the one in ref [21]. For the bulk samples, since our model implements multi-region sampling, we first evaluate the multivariate VAF distribution (which is a joint probability distribution of all sampled bulk VAFs) and then calculated the Euclidean distance between the obtained empirical probability distribution vectors:

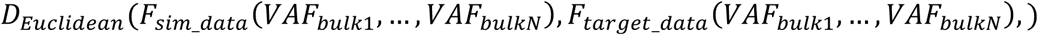

With single cell samples, we constructed phylogenetic trees per tumour and used different tree-based summary statistics to evaluate the distance. Since the inferred phylogenetic tree branch length is proportional to the number of unique mutations belonging to a node, we decided to compare the vectors of all branch lengths (between a simulated and target tumour trees) by computing the Wasserstein distance. For the subclone introduction time *t*, death rate *d* and the boundary driven growth parameter *a*, we chose to compare the vectors of branching times for each node of the phylogenetic trees.

Since our spatial simulation model requires more computation time and space to generate data samples close to a realistic tumour sizes, we are limited to run the ABC framework with a small tumour size (∼100k cells) or simulate smaller amount of datasets per inference, both of which can significantly affect the outcome. To therefore speed up our ABC framework we implemented a Sequential Monte Carlo (SMC) algorithm to increase the acceptance rate of the simple ABC rejection algorithm. Our ABC SMC algorithm uses sequential importance sampling by running several rounds of resampling around the accepted parameters (correlating the rounds), and gradually decreasing the acceptance threshold while converging to the posterior distribution. This approach significantly increases the acceptance rate of the simulated datasets [44].

Our implementation of the ABC SMC algorithm is as follows:

1. Initialise the indicator to rounds *r* and the acceptance threshold *ε*
2. ***If*** *r* = 1
  2.1. Run the simple ABC rejection algorithm (described above).
  2.2. Order the simulated parameters set according to their corresponding distance values.
  2.3. Keep the top Q per cent of the parameters.
3. ***Else***
  3.1. Sample next particle *θ* = (*u, t, s, d, a*) from the accepted set of parameters from round *r* − 1 with weights *W*_r-1_.
  3.2. <u>P</u>erturb each sampled parameter *p*_*i*_ using uniform perturbation kernel
  *K* = *Unif*(*p*_*i*_ − *σ, p_i_* + *σ*), where 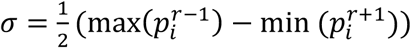.
  3.3. ***<u>If</u>*** *<u>π</u>*(*<u>θ</u>*<u>) > 0</u>, keep *θ*
  ***Else***<u>go to step 3.2.</u>
  3.4. <u>Simulate</u> data from the model using the sampled particle *θ*.
  3.5. Calculate distance D between the target and the simulated data.
  3.6. ***If*** *D* < *ε*, keep *θ*
  ***Else***<u>go to step 3.1.</u>
4. Calculate the weights for all accepted particles 1 ≤ *j* ≤ *N*:
  4.1. ***If*** *r* = 1, set *W*_(j,r)_ = 1
  4.2. **Else**

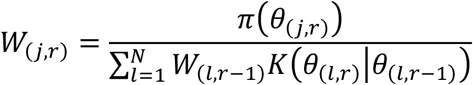
5. Update the threshold *ε* to the top Q-th percentile of the accepted particles.
6. Repeat until *ε* is less than a desired convergence threshold.

Our ABC-SMC framework tries to recover all the parameters (referred as a particle in the algorithm above) at the same time. We notice that once one of the parameters converges, the acceptance rate decreases significantly. As a workaround for this, we decided to fix the converged parameter at the inferred value (mode of its posterior) and rerun the inference varying the rest of the parameters until other parameters converge, and repeat the procedure.

### Phylogenetic tree reconstruction

For Figure 4 and parameter inference framework with single cell sequenced trees we used maximum parsimony algorithm implemented in paup [ref] phylogenetic tree inference software. For the genotyped phylogenetic trees in Figure 4, we manually constructed input genotype files for paup, by recording presence/absence of a given mutation from the sampled 200 cells with respect to the reference mutations list (in our case mutations list taken from a bulk sample).

### Neutrality test

To test for the presence of selection and the mutation rate inference, we fit 1/f distribution to the empirical cumulative distributions of sampled VAFs using the R package developed in ref [20].

## Acknowledgments

This work has been supported by Medical Research Council funding to A.S. (MR/P000789/1). A.S. is also supported by the Wellcome Trust (202778/B/16/Z) and Cancer Research UK (A22909). T.G. is supported by the Wellcome Trust (202778/Z/16/Z) and Cancer Research UK (A19771). We acknowledge funding from the National Institute of Health (NCI U54 CA217376) to A.S and T.A.G. This work was also supported a Wellcome Trust award to the Centre for Evolution and Cancer (105104/Z/14/Z).

## Supplementary Figures

**Figure S1.**
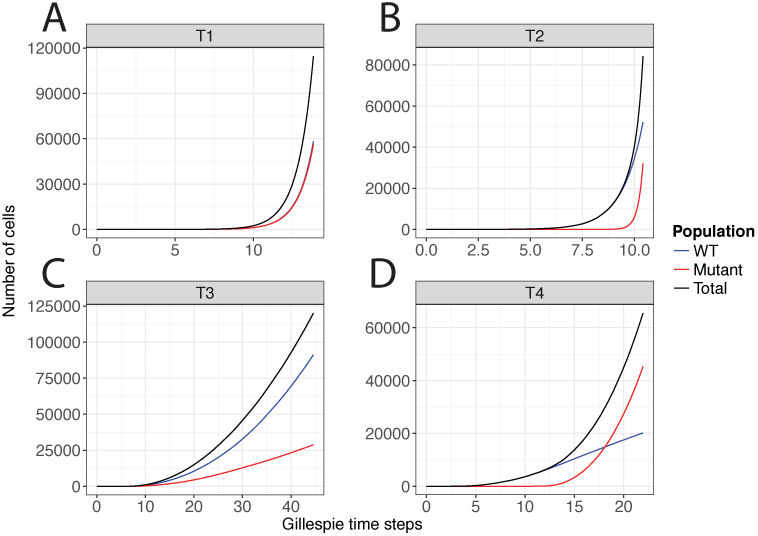
Growth curves. Tumour cell population growth curves for each of the representative cases: **(A)** neutral homogeneous, **(B)** selective homogeneous, **(C)** neutral boundary driven, **(D)** selective boundary driven. Wild type (WT) and mutant growth curves are plotted separately in addition to the whole population growth curves. Without the spatial constraints of our model, the growth curves are exponential as expected. **(A, B)** With the boundary driven growth the growth becomes polynomial. We can also see for the tumours with selection **(B, D)** how the mutant subpopulation outcompetes wild type cell population.

**Figure S2.**
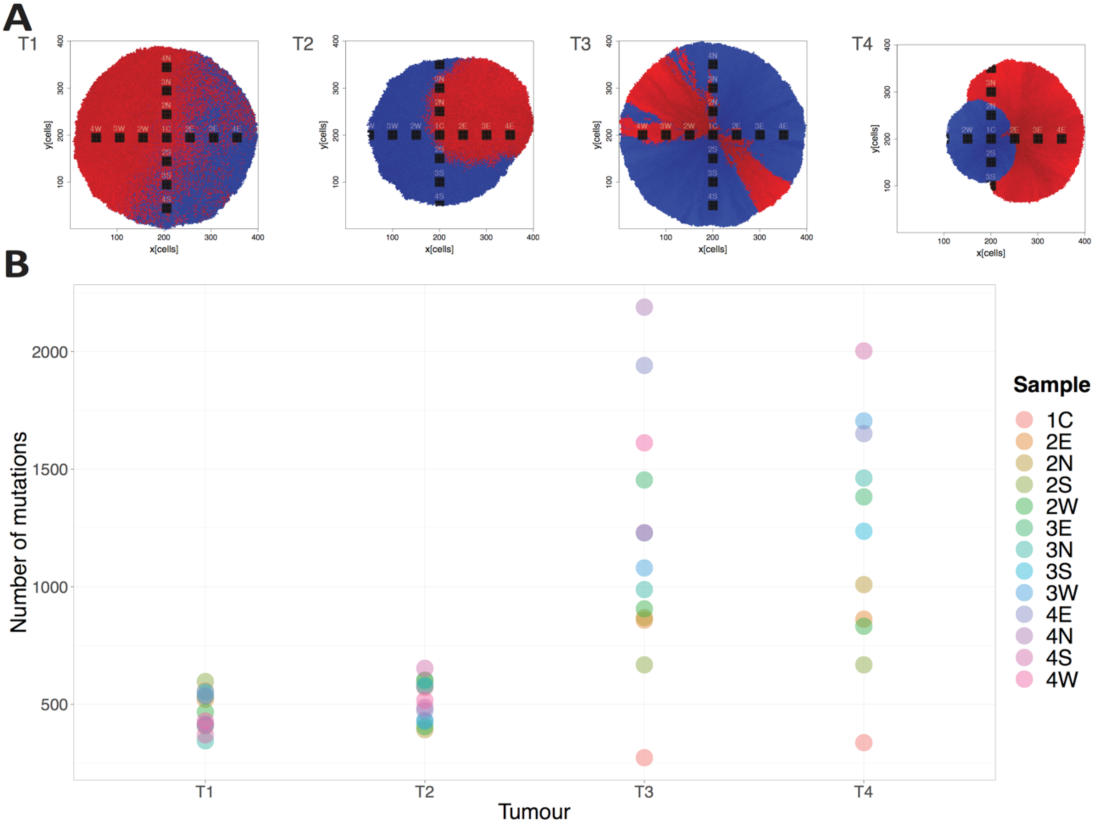
Mutational load comparison for different growth cases. **(A)** We sample each representative example tumours (T1 – neutral homogenous, T2 – selective homogenous, T3 – neutral boundary driven, T4 – selective boundary driven) from the tumour centre (bulk sample C1) towards the periphery following the concentric circles in four directions: W – west, E – east, N – north, S – south. The bulk indexes (2W, 3W, 4W) are proportional to the distance from the centre to the periphery. **(B)** We observe how number of mutations per bulk sample increases proportionally to the distance from the tumour centre to the sample location. Also, the total number of mutations are much higher for the constrained boundary driven growth than for the homogenous tumour due to increased cell turnover in the former case.

**Figure S3.**
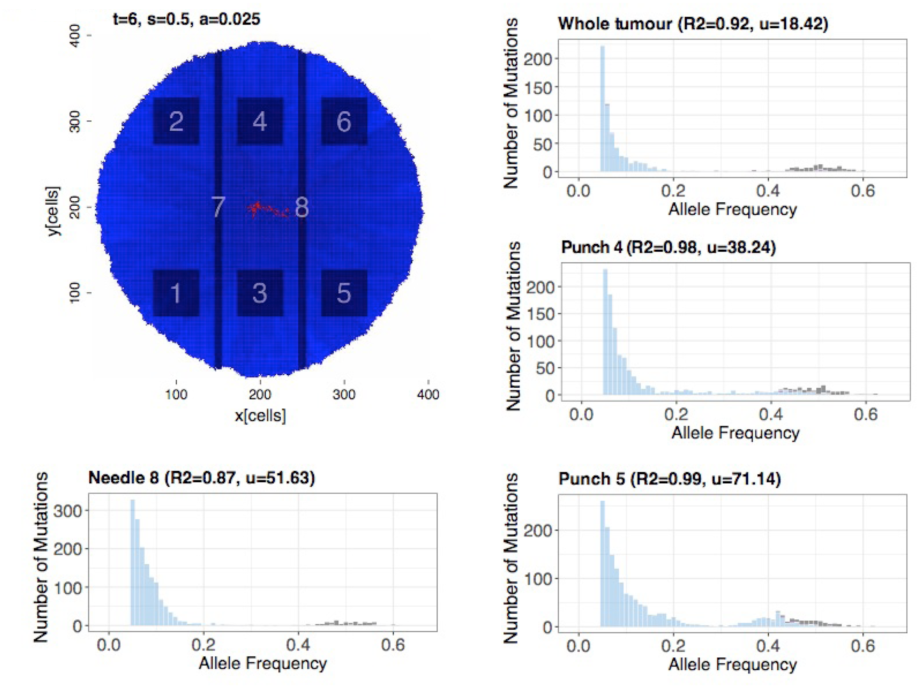
Example of imprisonment. Example of selective boundary driven growth when the mutant subpopulation gets trapped within the wild type population.

**Figure S4.**
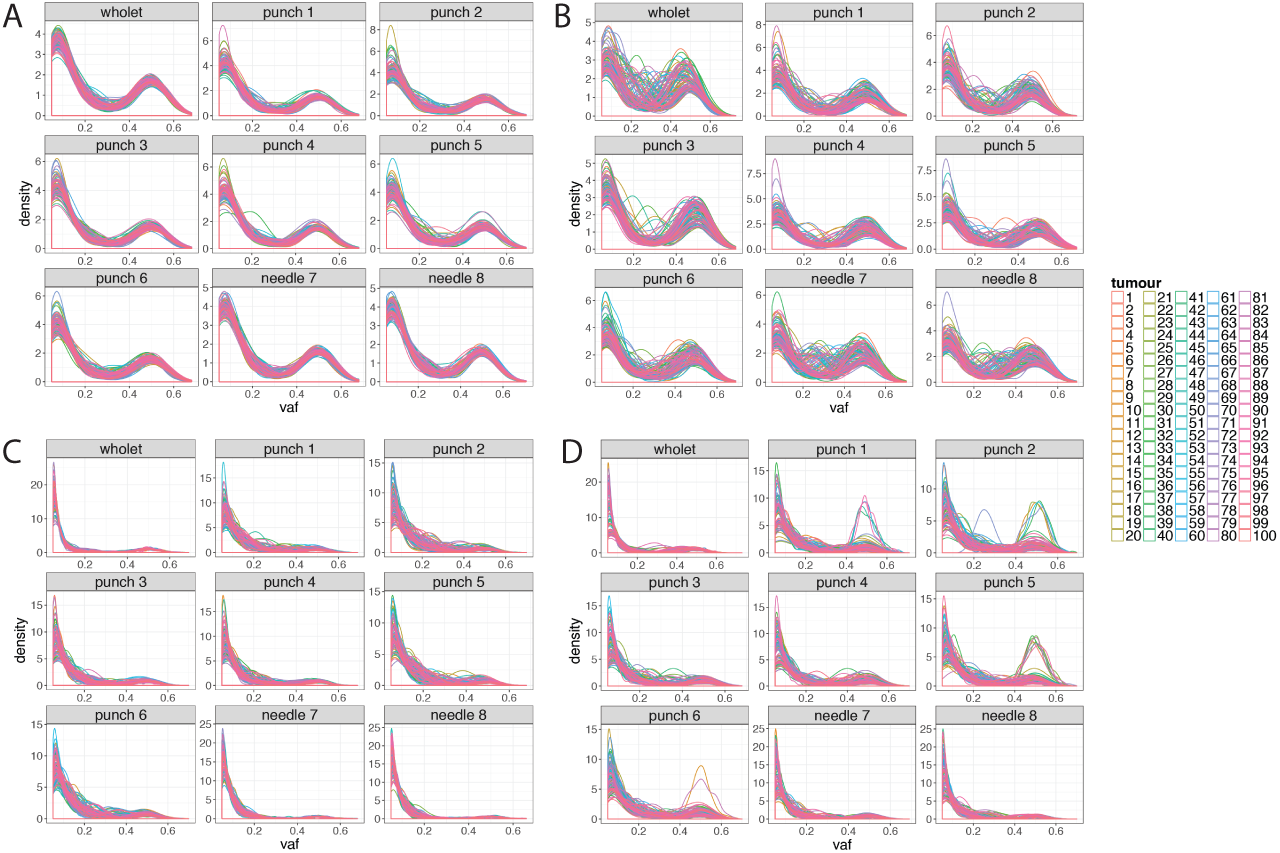
The effect of stochasticity and sampling bias on the shapes of VAF distributions for the four representative scenarios from Figure 2. For each of the representative cases: **(A)** neutral homogeneous, **(B)** selective homogeneous, **(C)** neutral boundary driven, **(D)** selective boundary driven, we simulated 100 different runs of each case keeping the underlying parameters constant and varying only the random seed of the simulation. For each simulated tumour, we constructed needle and punch biopsy sample VAF distributions along with the whole tumour VAFs. Overall there is a less variation among the distributions for neutral **(A,C)** versus selective **(B,D)** cases. In addition, punch biopsy VAFs scatter more than needle biopsy samples in comparison to the whole tumour VAF distributions.

**Figure S5.**
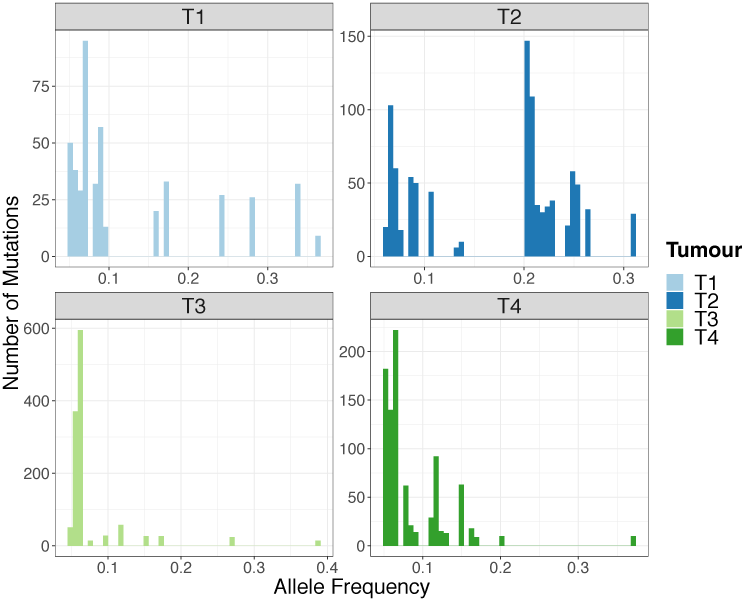
Allele frequency distributions derived from single cell sequencing. *We construct the allele frequency distributions from sequencing the randomly sampled 400 single cells (same as in Figure 4) from the four representative tumour examples: T1 – neutral homogenous, T2 – selective homogenous, T3 – neutral boundary driven, T4 – selective boundary driven.*

**Figure S6.**
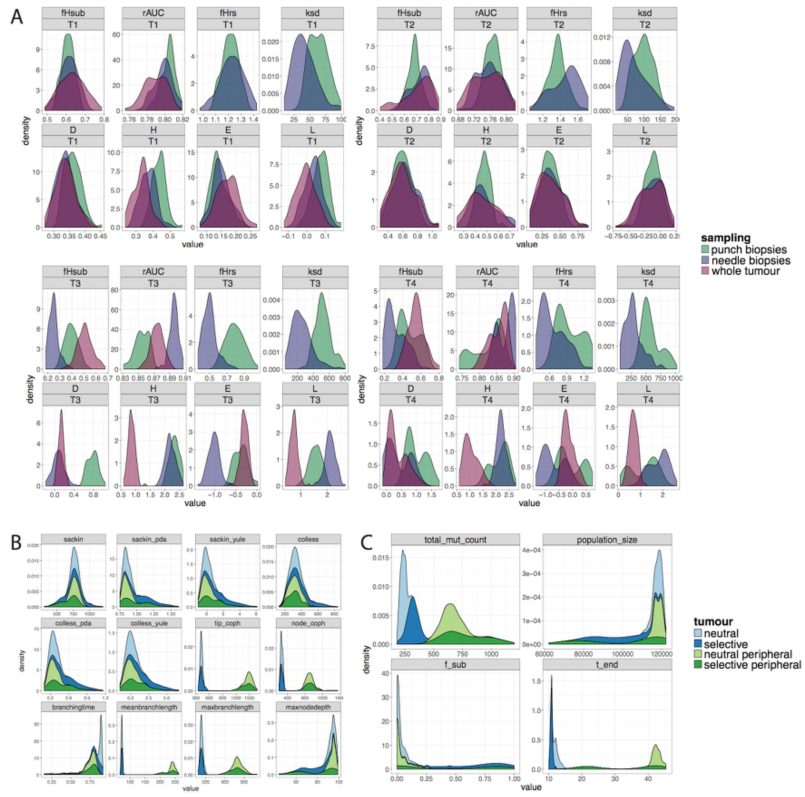
Comparing site frequency spectrum and phylogenetic tree balance index statistics for each representative scenario and sampling strategy. **(A)** Distributions of Site Frequency Spectrum (SFS) based summary statistics (discussed in the materials and methods section) for 100 different runs of the four cancer evolution scenarios: T1-neutral homogeneous, T2 - selective homogeneous, T3 - neutral boundary driven, T4 - selective boundary driven. For the homogenous tumour cases, we see the distributions from most of the summary statistics from needle biopsy samples are closer to whole tumour shapes than punch biopsy examples. Although, overall there is not as much variation among the distributions for all the three sampling strategies compared to the same distributions from the boundary driven tumour cases. **(B)** Distributions of different summary statistics (materials and methods section) from single cell sampling (100x) phylogenetic trees for the four representative cases. The balance index based statistics (sackin, colless with their different normalisation approaches – Yule, PDA) seem to have similar shapes among all four tumour cases, while tip and node Cophenetic distance based statistics show different trends for neutral versus selective examples with not observable variation between homogenous and boundary driven tumours. Branch length based statistics give similar results as cophenetic distances. Only one statistics, maximum node depth, tend to have longer flat tails for boundary driven tumours compared to homogenous tumour simulations. **(C)** For each of four tumour examples, we compare total number of passenger mutations and final population sizes along with the time the simulations finish and the final frequency of the new sub-population (introduced after a driver event). There is a higher number of mutations accumulated for the tumours growing on the periphery compared to the homogenous tumour simulations due to high cell turnover of the former cases. The subclonal frequencies for the neutral cases peak near zero possibly indicating to the strong effect of random drift. For the selective cases, f_sub distributions are more spread out indicating again the higher effect of stochasticity for these tumours as well as random drift.

**Figure S7.**
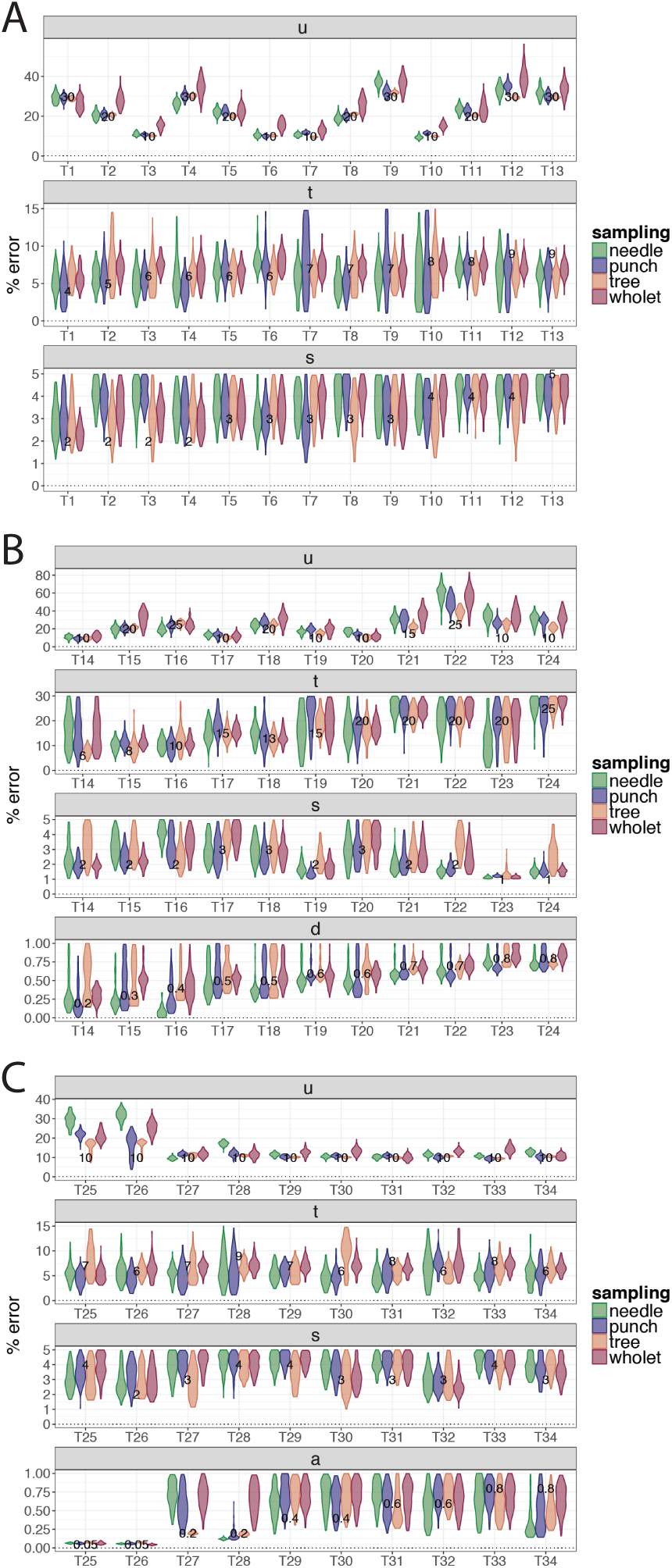
Posterior distributions of model parameters from each synthetic tumour. The violin plots of the posterior distributions for each model parameter per synthetic tumour inferred by our ABC-SMC framework. The three sets of tumours corresponding to the three tumour growth scenarios are plotted separately: exponential **(A)**, death **(B)** and boundary driven **(C)**.

**Figure S8.**
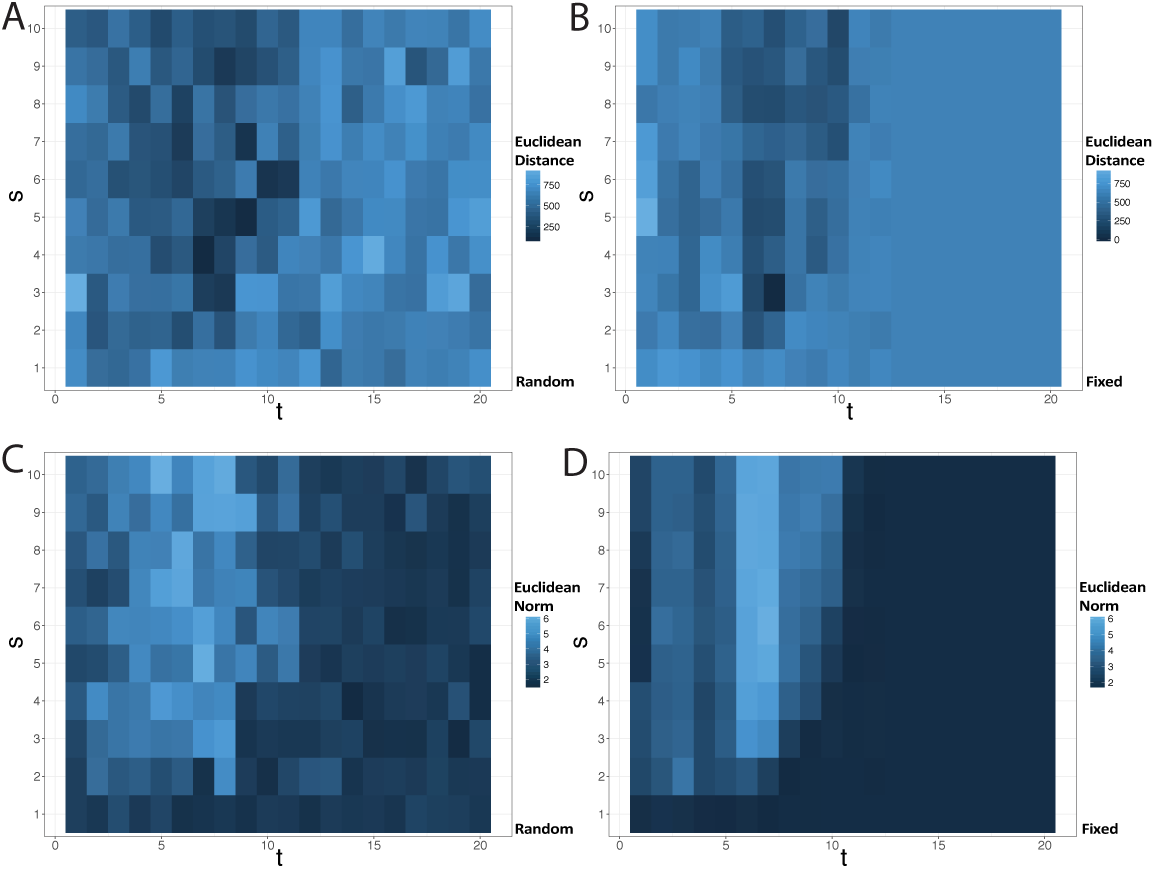
The effect of stochasticity on the dependence of t and s parameter combinations on the VAF distribution. To explore the interdependence of the parameter pair t and s, for their different values we simulate tumour growth while fixing all the other parameters (2D grid size=400, u=10, d=0, a=1). We summarised the obtained tumours by calculating either the Euclidean norm of the obtained whole tumour VAFs **(C, D)** or the calculating Euclidean distance between the cumulative VAF distributions of the simulated and a chosen target tumour (in this case target tumour parameters are t=7, s=3) **(A, B).** To reduce the effect of stochasticity we fix the random seed in **(B)** and **(D)** and they indeed showed less scattered patterns of **(A)** and **(C)** plots respectively. We observe how the closest tumours to the target tumour are the ones with the similar parameter values, although the effect of stochasticity is still high as a different t and s parameter combinations can give similar VAF statistics. We also see how low t or high s values result in higher VAF Euclidean norms indicating to higher number high frequency mutations.

